# APOL1 G1 and G2 risk alleles modulate severity of diet-induced obesity in a transgenic mouse model

**DOI:** 10.64898/2025.12.23.696293

**Authors:** Andrew O. Kearney, Johnson Y. Yang, Esther Liu, Matthew Wright, Margaret Chen, Jiayi Kong, Grant Barish, Edward B Thorp, Jennie Lin

## Abstract

*APOL1* G1 and G2 risk alleles are associated with an increased risk of chronic kidney disease. However, a causal relationship between these alleles and cardiometabolic traits has not been experimentally validated. To address this gap, we placed transgenic *APOL1* G0, G1, and G2 FVB/NJ mice on a high-fat diet and analyzed them for weight gain as well as obesity-related cardiometabolic phenotypes. To test whether *APOL1* risk alleles modulate the inflammatory basis of obesity, we also exposed bone marrow derived macrophages from these mice to pro-inflammatory, pro-hypertensive, and dyslipidemic conditions.

*APOL1* high-risk allele female mice gained fat mass more readily than their low-risk female counterparts, while *APOL1* high-risk male mice gained fat mass less readily than low-risk males. A parallel sex difference was seen in expression of higher levels of *Abca1*, *Hmox1*, and *Srebf1* in lipid-loaded female bone-marrow derived macrophages expressing G1 and G2 APOL1, along with minor differences in cardiac function. However, this finding occurred independently of hypertension and insulin resistance, and with only minor albuminuria. Thus, our results highlight the importance of sex as a biological variable in future *APOL1* experiments.

## INTRODUCTION

African Americans are disproportionately affected by type II diabetes, hypertension, cardiovascular disease, and other related metabolic conditions such as obesity [1–5]. The burden of obesity falls most heavily on African American women, with African American men having rates of obesity comparable to other ancestral groups [6]. Given the close association between obesity and the development of other cardiometabolic conditions [7], elucidating the factors driving higher rates of obesity in African American women is critical for closing health gaps and disparities. For complex cardiometabolic traits, genetics has emerged as a major biological risk factor, and genome-wide association studies (GWAS) have determined that obesity is indeed a highly polygenic trait [8]. However, African-ancestry individuals are severely underrepresented in most obesity GWAS cohorts, which fail to capture the genetic diversity of African-ancestry populations [2].

African Americans are also at higher risk of developing chronic kidney disease (CKD) and experience a more rapid decline in kidney function after diagnosis [9, 10]. Unlike for obesity, multiple studies have identified genetic risk factors underlying this disparity, a portion of which is attributed to DNA variants in the gene *APOL1* encoding Apolipoprotein A1 (APOL1) [2, 11–14]. APOL1 is a trypanolytic factor providing protection against infection with *Trypanosoma brucei,* the causative agent of African Sleeping Sickness [15–17]. However, the subspecies *T. brucei rhodesiense* has gained resistance against APOL1 toxicity through the evolution of a serum resistance associated factor (SRA), which binds to and neutralizes APOL1. In response, two *APOL1* coding variants termed G1 (S342G and I384M) and G2 (Δ388N389Y) evolved in West Africa, both of which restore trypanolytic activity against *T. brucei rhodesiense* [18, 19]. As most African Americans have West African ancestry, *APOL1* G1 and G2 alleles are common in African Americans, with allelic frequencies of 23% and 13% respectively [18]. However, possessing two *APOL1* high-risk alleles is strongly associated with kidney disease through gain-of-function toxicity [11, 14, 18, 20–23].

Obesity, cardiometabolic disease, and CKD are often co-morbid: many patients with APOL1-mediated kidney disease may also be at higher risk for obesity and cardiometabolic disease. Given the high prevalence of *APOL1* high-risk alleles among African Americans, it has been hypothesized that APOL1 is involved in the development or progression of cardiometabolic diseases in patients of African ancestry. Supporting this hypothesis, APOL1 is expressed in many tissues and cell types outside of the kidney [24, 25], with particular attention focused on endothelial cells and immune cells. In endothelial cells (EC), *APOL1* high-risk alleles have been associated with impaired mitochondrial respiration, alterations in inflammation and membrane permeability, and increased risk of sepsis [26, 27]. Relevant to immune cell function, *APOL1* high-risk alleles alter cholesterol handling in macrophages, leading to the promotion of pro-atherosclerotic foam cell formation [28]. In addition, *APOL1* high-risk alleles kidney donor recipients have an increased risk of T-cell mediated rejection, regardless of donor genotype. In these patients, enhanced APOL1 expression was noted in activated CD4+/CD8+ T-cells [29]. A pro-inflammatory state is a hallmark of obesity [30], suggesting a possible interplay between APOL1 and obesity through immune perturbation.

Current evidence of APOL1’s role in cardiometabolic disease in humans is limited and conflicting, however. Many studies have found *APOL1* high-risk alleles to be unassociated or only weakly associated with adverse cardiovascular events [31–33]. Conversely, Ito et al. (2014) [34] found an approximate doubling of atherosclerotic cardiovascular disease risk among high-risk individuals, and Mukamal et al. (2015) [35] identified *APOL1* high-risk alleles as accounting for the majority of the racial disparity in peripheral artery disease and myocardial infarction. Nadkarni et al. (2021) [36] found that each *APOL1* high-risk allele increases BMI by an average of 0.36 kg/m^2^. These discrepancies most likely originate from the differences in study design, or as a result of worsening kidney function seen in high-risk *APOL1* individuals [32].

Despite the possibility of APOL1 exacerbating cardiometabolic conditions, no in-vivo experimental studies have been conducted to investigate the direct role of APOL1 in obesity. As such, we aimed to leverage a transgenic *APOL1* mouse model by subjecting the animals to a high-fat diet (HFD) to cause diet-induced obesity (DIO). We sought to experimentally demonstrate whether having two high-risk *APOL1* alleles (HR) increases susceptibility to DIO and associated cardiometabolic conditions as opposed to two low-risk *APOL1* alleles (LR). We also employed bone-marrow derived macrophages (BMDMs) and iPSC-derived endothelial cells (ECs) to investigate the molecular pathways elevated in LR and HR individuals driving these differences. As will be discussed, neither method satisfactorily emulated the desired phenotypes, limiting the scope of this investigation. However, we believe the limited findings presented do provide important considerations about the design of future APOL1-focused experiments in humans and animals.

## MATERIALS AND METHODS

### Animal Care

G0/G0-*APOL1* (G0), G1/G1-*APOL1* (G1), and G2/G2-*APOL1* (G2) transgenic FVB/NJ mice [37] were obtained from the Mutant Mouse Resource & Research Centers (University of California Davis, Davis, CA). Animals were housed in groups of three in ventilated cages under a 12-hour day-night cycle with constant humidity, temperature, and free access to water and the appropriate chow. Animal care and experimental procedures were conducted in accordance with the *Guide for the Care and Use of Laboratory Animals* established by the National Research Council and were approved by the Northwestern University Institutional Animal Care and Use Committee.

### Determination of APOL1 Protein Expression

Inguinal white adipose tissue (iWAT) was collected from 6-week-old G0, G1, and G2 mice (n = 2) and snap-frozen in liquid nitrogen. The tissue was homogenized in RIPA buffer (Sigma, St. Louis MO) with 1% protease-inhibitor cocktail (Thermo Fisher Scientific Inc., Waltham MA) using a Beadbug Homogenizer (Benchmark Scientific, Inc., Sayreville NJ). Equal amounts (25 µg) of total protein were separated by electrophoresis on 4-12% gradient agarose gel (Thermo Fisher Scientific Inc., Waltham MA) and transferred to a polyvinylidene difluoride membrane using an iBlot2 dry transfer system (Thermo Fisher Scientific Inc., Waltham MA). Membranes were blocked with 10% superblock solution (Thermo Fisher Scientific Inc., Waltham MA) and probed with anti-APOL1 antibody (Genentech, South San Francisco CA) at 4°C overnight. Blots were washed and probed with anti-mouse HRP-conjugated secondary antibody (Abcam, Cambridge UK) for 2 hours at ambient temperature. Blots were detected by ECL solution (Thermo Fisher Scientific Inc., Waltham MA) and imaged on an iBright 1500 system (Thermo Fisher Scientific Inc., Waltham MA). Blots were stripped with stripping buffer (15g glycine, 1g SDS, 10 mL Tween 20 / 1L H2O) and re-blocked as described above. Blots were incubated with rabbit anti-ß-actin antibody (Cell Signaling Technology, Inc., Danvers MA) overnight at 4°C, then continued to development as described above using Goat anti-Rabbit HRP-conjugated secondary antibody (Abcam, Cambridge UK). Protein levels were determined using iBright image analysis software and are expressed relative to ß-actin levels.

### High-Fat Diet

Two independent diet-manipulation experiments were conducted. In the first, animals were enrolled at 6 weeks of age according to their sex (M or F) and genotype (G0, G1, or G2) and randomly assigned to the control diet (16 weeks of 10 kcal% low-fat diet (LFD)) or the weight gain diet (>60 kcal% HFD) for 16 weeks (Research Diets Inc., New Brunswick, NJ) (n = 6; 72 total). Animals were weighed three times per week for the first 10 weeks, after which they were weighed once weekly. Fasting serum and urine were collected at baseline and endpoint. After a 6-hour fasting period, whole blood was collected via a retro-orbital method under isoflurane anesthesia, followed by separation via centrifugation. Urine was collected by massaging the bladder of the animal. Body composition was assessed non-invasively at week 16 using an EchoMRI-E26-388-MT system (EchoMRI, Houston TX). Blood pressure was measured at 16 weeks as described below. Animals were euthanized at the end of the 16-week period via thoracotomy and cardiac perfusion with PBS while under ketamine/xylazine anesthesia. Inguinal white adipose tissue, epidydimal white adipose tissue, partial liver, whole heart, and whole kidney tissue was collected and either fixed in 10% formalin solution for 24hr at 4°C or snap-frozen in liquid nitrogen.

In the second experiment, animals were enrolled at 8 weeks of age according to their sex (M or F) and genotype (G0, G1, or G2) and randomly assigned to the control diet (C; 24 weeks of LFD), the weight gain diet (WG; 8 weeks of LFD then 16 weeks of HFD), or the weight loss diet (WL; 16 weeks of HFD then 8 weeks of LFD). Body weight was measured once every two weeks. Fasting serum, urine, and body composition were collected at baseline, week 8, and week 16 as described above. Additionally, mice had cardiac parameters measured by echocardiogram and glucose tolerance assessed by oral glucose tolerance testing (oGTT) at week 23, both described below. Fasted serum, urine, blood pressure, and tissues were collected at week 24 as described above.

### Blood Pressure Evaluation

Systolic, diastolic, and mean arterial blood pressure (MAP) was assessed at week 16 via the tail cuff method using a CODA High Throughput System (Kent Scientific, Torrington CT). Animals underwent three acclimation sessions over at least three days prior to collecting blood pressure measurements. Blood pressure was collected between the hours of 9:00am-12:00pm to minimize fluctuations due to circadian cycles. Animals were warmed during the collection period using an external heating pad. At least thirty collection cycles were collected per animal. Collection cycles with <10 µL of total flow were rejected, and the remaining measurements were averaged into a single value.

### Measurement of Albuminuria

Albuminuria was determined by calculating the urinary albumin to creatinine ratio (uACR) according to the following formula: [uACR = Urine albumin (µg/mL) × Urine Creatinine (mg/dL) / 100]. Urine albumin levels were determined using the Exocell Albuwell M ELISA kit, and urine creatinine levels were determined using the Exocell Creatinine Companion Kit (Ethos Bioscience, Logan Township NJ).

### Profiling of Metabolic Analytes

Circulating levels of IL-6, MCP-1, TNF-α, Insulin, and Leptin in fasted blood serum were determined using the MILLIPLEX® Mouse Adipokine Magnetic Bead Panel – Endocrine Multiplex Assay (Millipore, Burlington MA) on a Luminex 200 Instrument System (Diasorin, Saluggia IT). Fasting glucose was determined using the Freestyle Lite Glucometer (Abbott, Chicago IL) and was used to calculate HOMA-IR according to the following formula: [HOMA-IR = fasting serum glucose (mg/dL) × fasting serum insulin (mU/L) / 405].

### RNA Isolation and Quantitative Real-Time qPCR

Total RNA was extracted using TRIzol (Thermo Fisher Scientific Inc., Waltham MA) following manufacturer’s instructions. Snap-frozen mouse tissues were homogenized using a Beadbug Benchtop Homogenizer using 1.5mm zirconium beads (Benchmark Scientific, Inc., Sayreville NJ) filled with TRIzol. Cell culture samples were homogenized using a cell scraper immersed in TRIzol. The total RNA was purified with the RNA Clean and Concentrator-25 kit (Zymo, Irvine CA) and quantified by Nanodrop. Reverse transcription was performed using High Capacity cDNA Reverse Transcription Kits (Thermo Fisher Scientific Inc., Waltham MA). Polymerase chain reaction analysis was performed with PowerUp® SYBRGreen PCR Master Mix (Thermo Fisher Scientific Inc., Waltham MA) on a QuantStudio™ 5 Real-Time PCR System (Thermo Fisher Scientific Inc., Waltham MA). PCR results using tissue-derived cDNA were normalized to mouse *Atpfa1* gene expression. PCR results using cell culture-derived cDNA were normalized to eukaryotic *18S* gene expression. Data was analyzed and presented as fold-change from control G0/G0 samples using the 2^−**ΔΔCt**^ method.

Real-time PCR cycler conditions were: 120 seconds at 50°C, 600 seconds at 95°C, followed by 40x cycles of [15 seconds of 95°C + 60 seconds at 60°C]. Temperature was ramped up and down at ±1.6 °C/s.

### Histology

Heart, kidney, liver, spleen, inguinal white adipose tissue, and epidydimal white adipose tissue were fixed in 10% formalin solution for 24 hours at 4°C. Tissues were washed three times with ice-cold PBS following long-term storage in 70% ethanol solution at ambient temperature.

Tissues were embedded in paraffin and cut into 4 µm-thick sections on positively-charged slides (Assure+, Creative Waste Solutions, Inc., Tualatin OR). Dewaxed sections were routinely stained with hematoxylin-eosin (H&E) dyes for histological evaluation. Kidney sections were also processed for chromogenic immunohistochemistry (IHC) for CD68 (dilution: 1:200; Cat# 97778S, Cell Signaling Technology, Danvers MA). iWAT sections were processed for CD206 (dilution: An anti-rabbit antibody-horseradish peroxidase (HRP) polymer conjugate (MACH2, Cat# MHRP520, Biocare Medical, Patheco CA) was used in conjunction with the chromogenic substrate 3,3’-diaminobenzidine (DAB) (Vector Labs, Newark CA) to visualize the primary antibody binding sites. IHC slides were counterstained with hematoxylin.

Three 500µm x 375µm areas were analyzed per kidney section: the superior cortex, inferior cortex, and lateral cortex. CD68 immunostaining was scored as area stained (permille) using the <u>IHC Toolbox plugin</u> (Version 1.0) in ImageJ (Version 1.54d) [38]. Using an automatic statistical color selection algorithm, DAB-H colored pixels are separated from background pixels. Pixels below a brightness of 65/255 were counted as positive pixels for the purpose of reducing noise.

Three 333.3µm x 250µm areas were analyzed per iWAT section. CD206 immunostaining was scored as described above. Cell surface area was estimated by multiplying the measured length by the measured width, approximating the cell’s surface as a rectangle. A minimum of 10 cells were measured per area (minimum of 30 cells per iWAT section).

### Oral Glucose Tolerance Testing

At week 23, food was removed from cages for a 16-hour overnight fasting period. Animals were administered 20% dextrose solution via oral gavage at a dose of 2g dextrose / 1kg lean body mass as determined by EchoMRI. Blood was collected prior to oral gavage as well as 15, 30, 45, 60, 90, and 120 minutes after gavage. Blood glucose was measured using Freestyle Lite Glucometer (Abbott, Chicago IL). Area-under-the-curve (AUC) was determined from the resulting oGTT plots using a trapezoidal approximation [39].

### Echocardiographic Image Acquisition

Transthoracic echocardiography was performed using established protocols with adaptations for both long- and short-axis views [40–42]. On the day of imaging, hair was removed from the left thoracic region using a depilatory cream (Nair) applied with cotton swabs to ensure complete clearance from the mid-chest to the axilla and neck.

Mice were anesthetized with 2.5–3% isoflurane in 0.5 L/min oxygen and positioned supine on a heated platform maintained at 37–38°C (FUJIFILM VisualSonics, Toronto, ON, Canada). Limbs were secured using medical tape, and isoflurane levels were gradually reduced to 1–2% to maintain physiologically stable and comparable heart rates during image acquisition.

Echocardiographic imaging was conducted using ultra-high-frequency linear array transducers—MX400 (18–38 MHz, center frequency 30 MHz, axial resolution 50 μm) for mice—connected to Vevo® 3100 imaging systems (FUJIFILM VisualSonics, Toronto, ON, Canada). For parasternal long-axis imaging, B-mode cine loops were acquired to visualize the left ventricle (LV) from apex to base at its maximal dimension. For short-axis imaging, the transducer was positioned at ∼80° to the mouse’s head-tail axis, with the head facing away from the operator. The imaging platform was tilted ∼30° to the left and ∼20° toward the operator to optimize visualization of the LV. Ultrasound gel was applied liberally to the chest, and the transducer was carefully lowered to make contact with the gel. Imaging proceeded from the base toward the apex until the posterior wall of the heart was visualized. Optimal cross-sectional views of the LV chamber were obtained by fine adjustments in probe angle and platform position, avoiding interference from papillary muscles. M-mode was employed to acquire four high-quality cardiac cycles in each view. All images were stored in raw DICOM format for offline analysis.

### BMDM Generation

At 8-12 weeks of age, G0, G1, and G2 mice were euthanized by CO_2_ inhalation and bone marrow was harvested from their femurs and tibias. The bone marrow was cultured in RPMI media supplemented with 10% heat-inactivated fetal bovine serum, 20% L929 cultured media, 1x GlutaMAX, and 1% penicillin/streptomycin (Gibco, Waltham MA). Cells were cultured in a 37°C incubator with 5% CO2. Media was replaced every 3 days until macrophages were used on days 7-10.

### OxLDL Loading

Mature BMDMs were pre-treated with 5 ng/mL of interferon gamma for 8 hours (PeproTech, Inc., Cranbury NJ). The cells were then washed 3 times with cold PBS, followed by 72-hour incubation with 5 ng/mL interferon gamma and/or 50 µg/mL OxLDL (Thermo Fisher Scientific Inc., Waltham MA). Cells were then washed and their RNA harvested as described above.

### Statistics

Statistical analyses were performed using GraphPad Prism 10.4.2. Two group comparisons were assessed using the Mann-Whitney U test unless otherwise specified. Multiple testing corrections was performed using the Two-stage step-up method of Benjamini, Krieger, and Yekutieli. Data plots are displayed as mean ± SEM. P-values <0.05 were deemed significant and indicated as *. P-values <0.01 were indicated as **. P-values < 0.001 were indicated as ***. P-values < 0.0001 were indicated as ****. For the purpose of statistical analysis, results from G1 and G2 mice were pooled together.

## RESULTS

### Alterations in Fat Accumulation are Sex-Dependent

Within human adipose tissue, *APOL1* is expressed in a variety of cells including adipocyte progenitors, endothelial mesothelial, and immune cells. While adipose tissue is known to express *APOL1*, adipocytes are minor contributors as compared to neighboring cell types (Figure 1A). Because *APOL1* is only expressed in humans, gorillas, and baboons, any non-human animal models require transgene insertion and proper expression. APOL1 protein expression in adipose tissue of transgenic APOL1 mice was confirmed with Western blotting (Figure 1B). Expression was robust with no significant difference between LR and HR mice.

**Figure 1.**
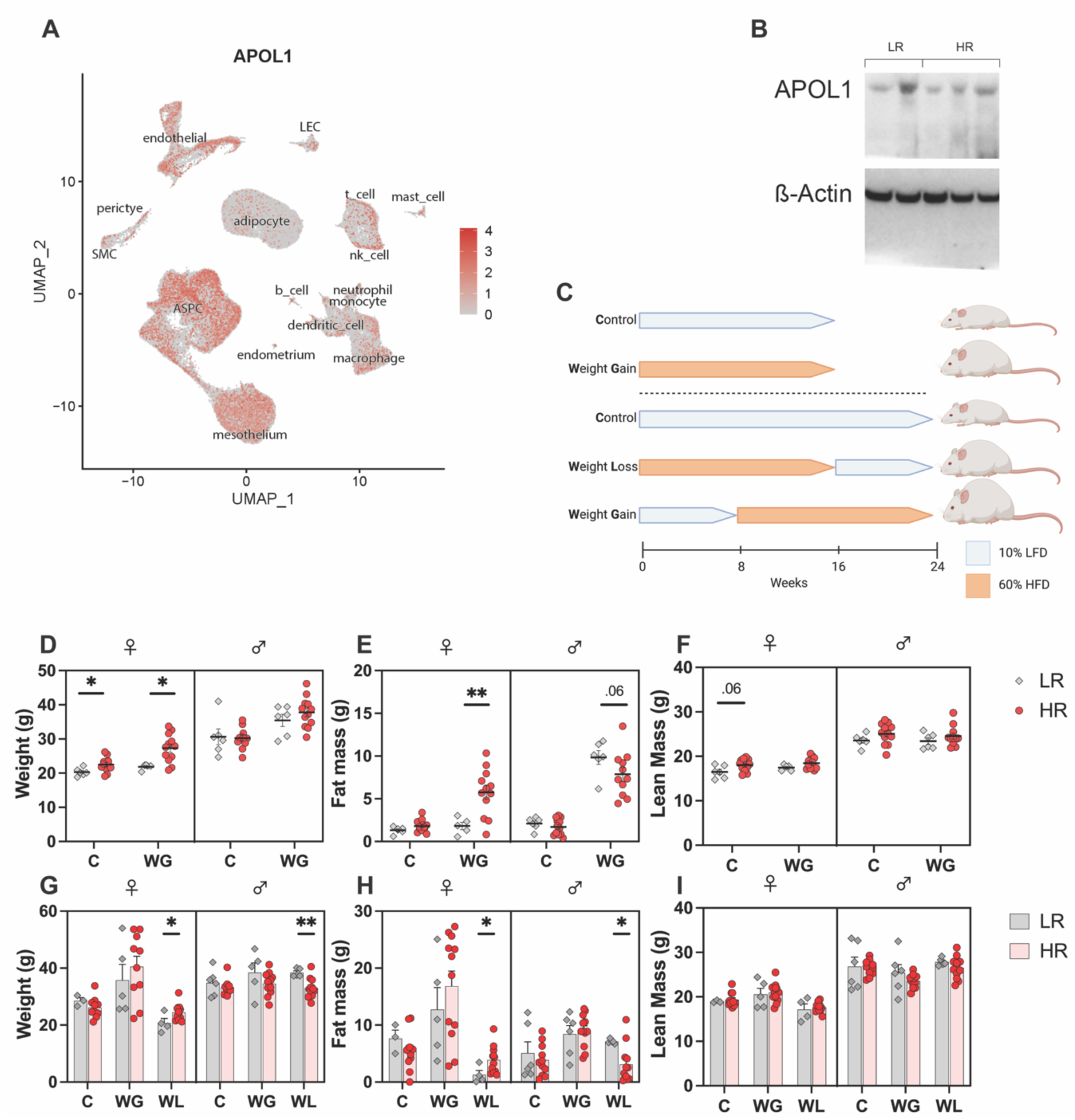
Experimental Setup and Gross Weight Results. **A)** Single-cell human gene expression of *APOL1*. UMAP feature plot of single cell RNA-seq of human white adipose tissue (PMID: 3529684). **B)** Anti-APOL1 and Anti-ß-Actin Western Blots of iWAT from G0, G1, and G2 mice at 8 weeks of age (n = 2). **C)** Schematic model of short-term (16-week) and long-term (24-week) diet experiments. Final **D)** gross weight; **E)** fat mass; and **F)** lean mass of short-term (16-week) experiment. Final **G)** gross weight; **H)** fat mass; and **I)** lean mass of long-term (24-week) experiment.

Two separate diet experiments were conducted (Figure 1C). The short-term experiment compared: 16 weeks of LFD, deemed control (C); and 16 weeks of HFD, deemed weight gain (WG). The groups contained equal numbers of both females and males, and equal numbers expressing G0, G1, and G2 *APOL1* alleles (n = 6). 16-weeks after diet initiation, HR females had greater gross body weight in both the C (+2.23 g, P = 0.026) and WG (+5.45 g, P = 0.018) groups (Figure 1D). NMR analysis revealed that LR and HR females had statistically similar lean masses (Figure 1F), while HR females had more fat mass only in the WG group (+3.93 g, P = 0.008) (Figure 1E). Conversely, WG HR males had less fat mass than WG LR males although this difference failed to reach significance (−3.04 g, p = 0.062) (Figure 1E).

The long-term experiment compared: 24 weeks of LFD, deemed control (C); 8 weeks of LFD followed by 16 weeks of HFD, deemed weight gain (WG); and 16 weeks of HFD followed by 8 weeks of LFD, deemed weight loss (WL). The WL cohort returned similar findings as the 16-week experiment. WL HR females had significantly higher total weight (+3.79g, p = 0.042) and fat mass (+2.63g, p = 0.030) at endpoint (Figures 1G-H), suggestive of more resistance to fat loss by caloric restriction in the HR genotype. WL HR males had significantly lower total weight (−5.36g, p = 0.008) and fat mass (−4.03g, p = 0.028) at endpoint (Figures 1G-H), which mirrored the trend suggested in the 16-week experiment. However, no significant difference in lean mass was observed in any cohort (Figure 1I), nor were statistically significant *APOL1* genotype differences noted outside of the WL group. WG HR females demonstrated a pattern suggestive of more weight gain compared to their LR counterparts but did not reach statistical significance. Implications of the differences in the 16 vs 24-week experiments will be discussed further.

In summary, these data suggest that the *APOL1* HR alleles alter the accumulation of fat in mice in a sex-specific manner. For female mice, HR increases the severity of obesity when fed HFD for 16 weeks, including when HFD is removed, mirroring potential differential responses to calorie-restricted diets. For male mice, HR unexpectedly decreases the severity of obesity.

### Increased Adiposity is Not Correlated with Insulin Resistance

We next investigated whether increased fat mass in HR females was reflected in surrogate measures of adiposity. The increased fat mass in the short-term WG HR females is reflected in the average fat cell size, which was approximately double that of short-term WG LR females (+1490 um^2, P = 0.022) (Figures 2A-B). Control HR females exhibited larger fat cells that control LR females as well (+551 µm^2, p = 0.048). There was no significant difference in fat cell size between LR and HR males.

**Figure 2.**
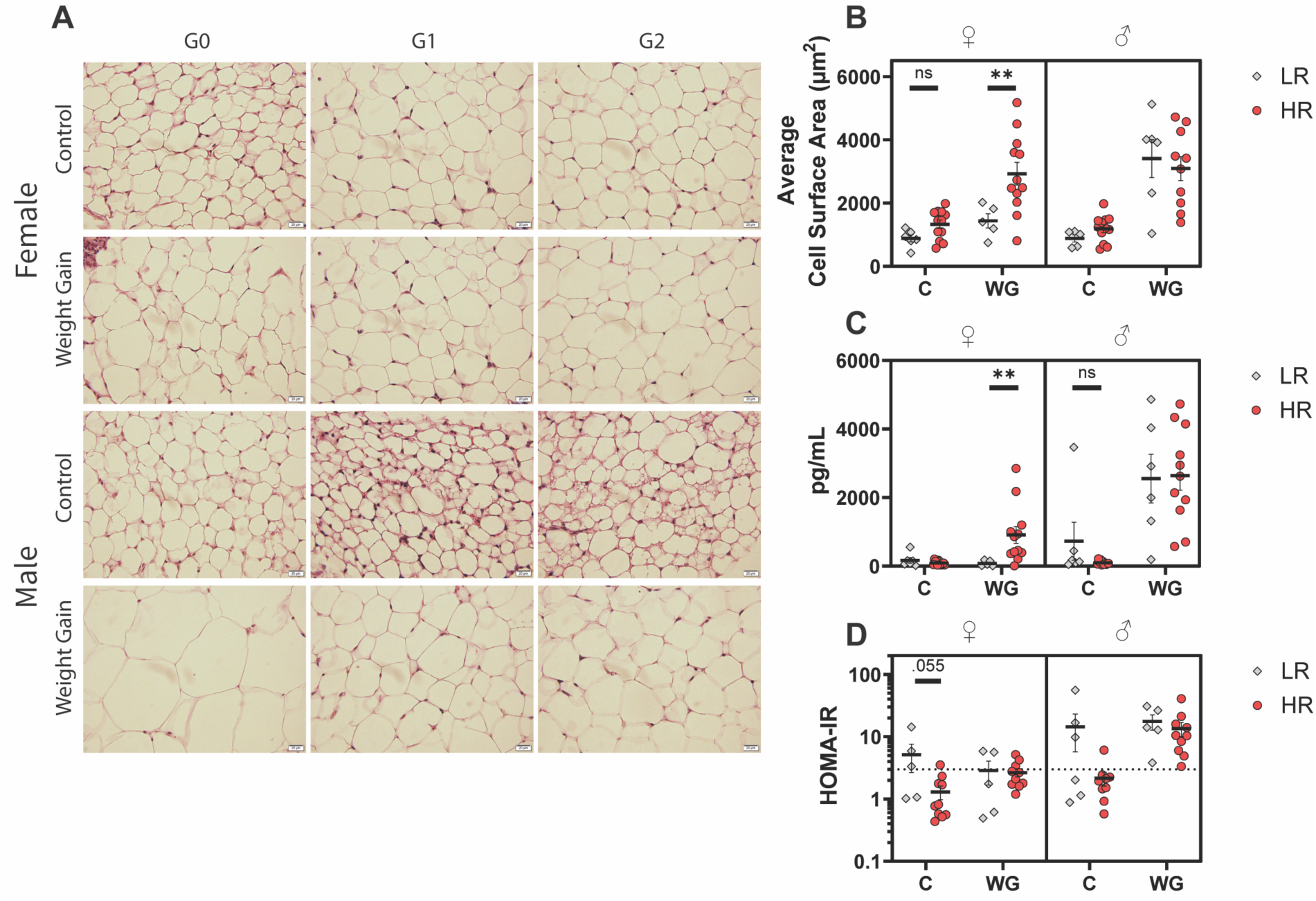
Adipose and Hormonal Responses. **A)** Representative images of H&E-stained iWAT tissue collected from short-term (16-week) experiment. Scale bar, 20µm. **B)** Quantification of cell surface area from H&E-stained iWAT sections. Cell surface area was estimated using a rectangular approximation. N = 30 cells / animal. **C)** Fasting serum leptin concentration of short-term experiment. Animals were fasted overnight for 16 hours prior to serum collection. **D)** Fasting HOMA-IR values of short-term experiment. HOMA-IR was calculated using fasting serum glucose and insulin concentrations according to the formula: [HOMA-IR = fasting serum glucose (mg/dL) × fasting serum insulin (mU/L) / 405]. H&E, hematoxylin and eosin; iWAT, inguinal white adipose tissue. HOMA-IR, homeostatic model assessment of insulin resistance.

Serum leptin levels similarly reflected the elevated fat mass in HR females only (+826.2 pg/mL, p = 0.048) (Figure 2C). Separate studies have shown an inconsistent link between fat mass and leptin levels in mice, with some groups reporting a near perfect linear relationship [43] and other groups reporting much weaker correlations [44]. A correlation is expected as leptin is secreted by adipocytes to promote satiety. We found the correlation to be relatively strong in both female (Leptin = 202.7*Fat – 243.3, R^2^ = 0.71) and male mice (Male: Leptin = 302.0*Fat – 140.3, R^2^ = 0.49), with the correlation stronger in females (Supplemental Figure 1E). Both metrics support our body composition findings and do not suggest obvious leptin dysregulation.

Paradoxically, control female and male mice appeared less glucose tolerant. For the short-term experiment, mean HOMA-IR values were elevated in C females (+3.82, p = 0.388) and highly elevated in control males (+12.27, p = 0.055) although neither reached significance (Figure 2D). For the long-term experiment, the area-under-the-curve of the oGTTs of control females (+4.585 µg*min/mL, p = 0.022) and control males (+6879 µg*min/mL, p = 0.003) were both significantly elevated (Supplemental Figure 6C). However, no significant difference in HOMA-IR or oGTT AUC was noted for the WL or either WG group. The lack of a significant decrease in glucose tolerance among WG mice is unexpected for a DIO model and suggests that HFD alone is incapable of inducing significant metabolic perturbation in this FVB/NJ APOL1 mouse model.

### APOL1 Genotype Differences Are Seen in Cardiac Strain

We next evaluated whether measurements relevant to cardiovascular health were impacted by *APOL1* genotype. Mean arterial pressure (MAP) did not differ significantly between LR and HR for any group in both the short-term and long-term experiments (Figure 3A). Of note, these diets were not high-salt. Urinary albumin:creatinine ratio (uACR), a marker of proteinuric kidney injury, was slightly elevated in WG HR females (+15.98, p = 0.007) and slightly decreased in WG HR males (−18.38, p = 0.031) at 16-wks, in a pattern similar to the risk-alleles differences in fat mass (Supplemental Figure 7C). However, these values are all within a relatively normal range of murine albuminuria, and do not represent significant injury.

**Figure 3.**
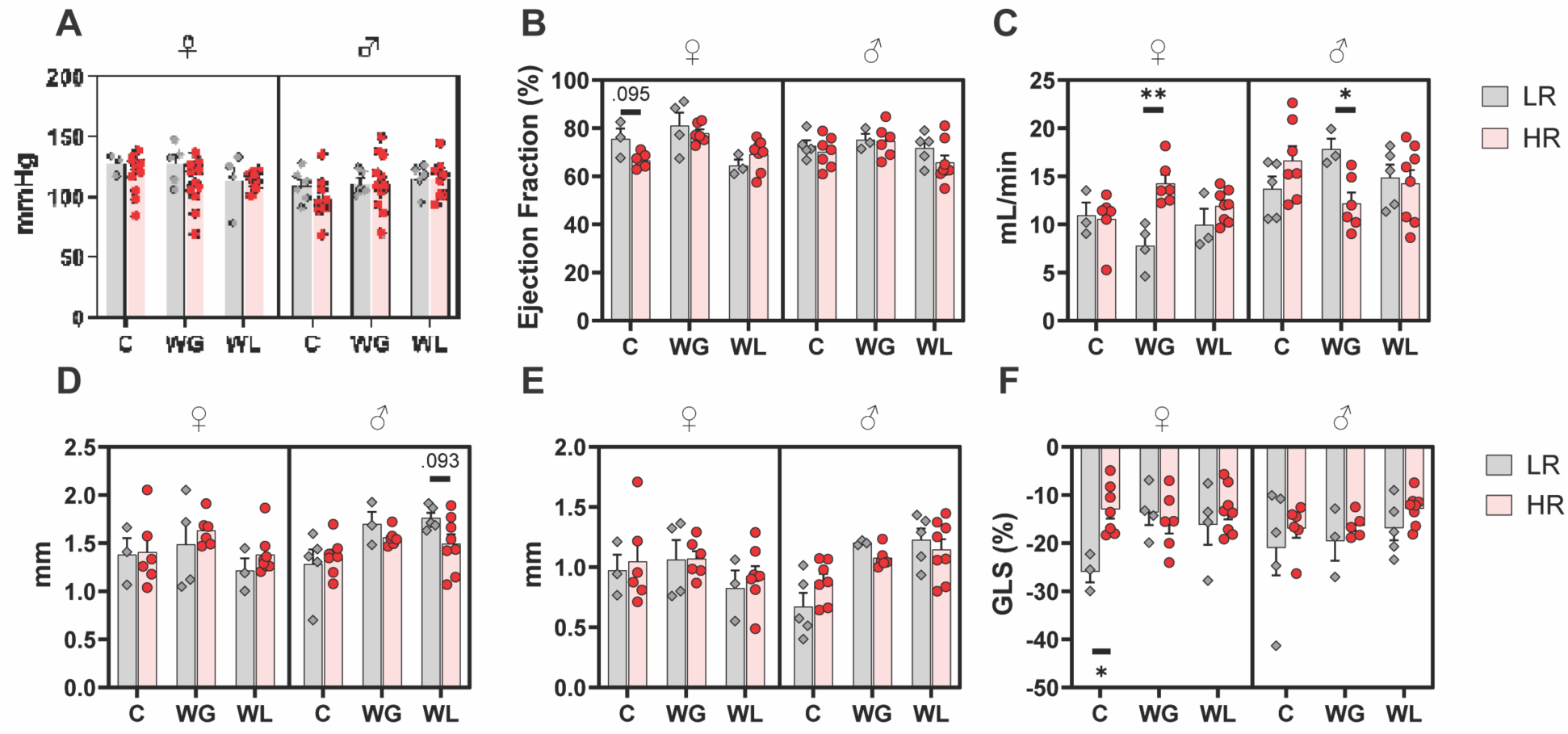
Genotype-specific Cardiac Strain Is Independent of Blood Pressure. **A)** MAP of long-term experiment collected via the tail-cuff method. **B)** EF(%) of long-term experiment. **C)** CO of long-term experiment. **D)** LVPW;s of long-term experiment. **E)** LVPW;d of long-term experiment. **F)** GLS% of long-term experiment. MAP, Mean Arterial Blood Pressure; EF(%), ejection fraction; CO, cardiac output; LVPW;s, left ventricular posterior wall end-systole; LVPW;d, left ventricular posterior wall end-diastole; GLS(%), global longitudinal strain.

The long-term mice underwent echocardiography at 23 weeks. Control LR females had slightly but not significantly elevated ejection fraction (+9.05%, p = 0.095) (Figure 3B) as well as a GLS% that was greater in magnitude (−12.98%, p = 0.017) (Figure 3F). Elevated ejection fraction and |GLS%| are both indicative of slightly better heart functioning in LR females on control diet, and this difference was observed in the absence of differences in blood pressure, suggesting underlying differences in cardiac mechanics independent of hypertension-related mechanisms. Of note, the genotype differences were abrogated by HFD. Left ventricular posterior wall thickness was slightly but not significantly depressed in WL HR males during systole (−.27mm, p = 0.093) although this difference was not apparent at end-diastole (Figure 3D-E). Cardiac output was elevated in WG HR females (+6.47 mL/min, p = 0.009) and depressed in WG HR males (−5.64 mL/min, p = 0.024) (Figure 3C). This pattern is similar to the pattern observed in gross weight and body fat, although the differences in cardiac output are seen in the WG rather than WL group. Alterations in cardiac function are subtle and non-pathogenic, but they do suggest that *APOL1* HR genotype may have affected both the functioning and structure of the heart. This is an important finding which warrants further investigation, given the association between *APOL1* genotype and poor cardiovascular outcomes that have been noted in human trials [34, 35]. We suspect that differences in cardiac function would be particularly apparent in a model of heart disease such as heart failure with preserved ejection fraction (HFpEF), and suggest future research focuses on the effect of murine *APOL1* genotype in HFpEF and related conditions.

### Sex Differences in APOL1 HR Macrophage Phenotype

Macrophage infiltration into adipose tissue is an important finding in obesity, and the chronic inflammatory signaling they trigger is thought to contribute heavily to co-morbidities [45]. In the absence of strong co-morbidities, we investigated phenotypic variation within macrophages derived from these mice. Staining of iWAT for CD206 showed a slight increase in CD206^+^ area in control HR females (+0.007 ‰, p = 0.048) and WG HR females (+0.019 ‰. p = 0.08), although only the former tested significant (Figure 4A-B). CD206 is a canonical marker of pro-resolving anti-inflammatory M2 macrophages [46], pointing towards a slight difference in inflammatory phenotype between LR and HR adipose macrophages.

**Figure 4.**
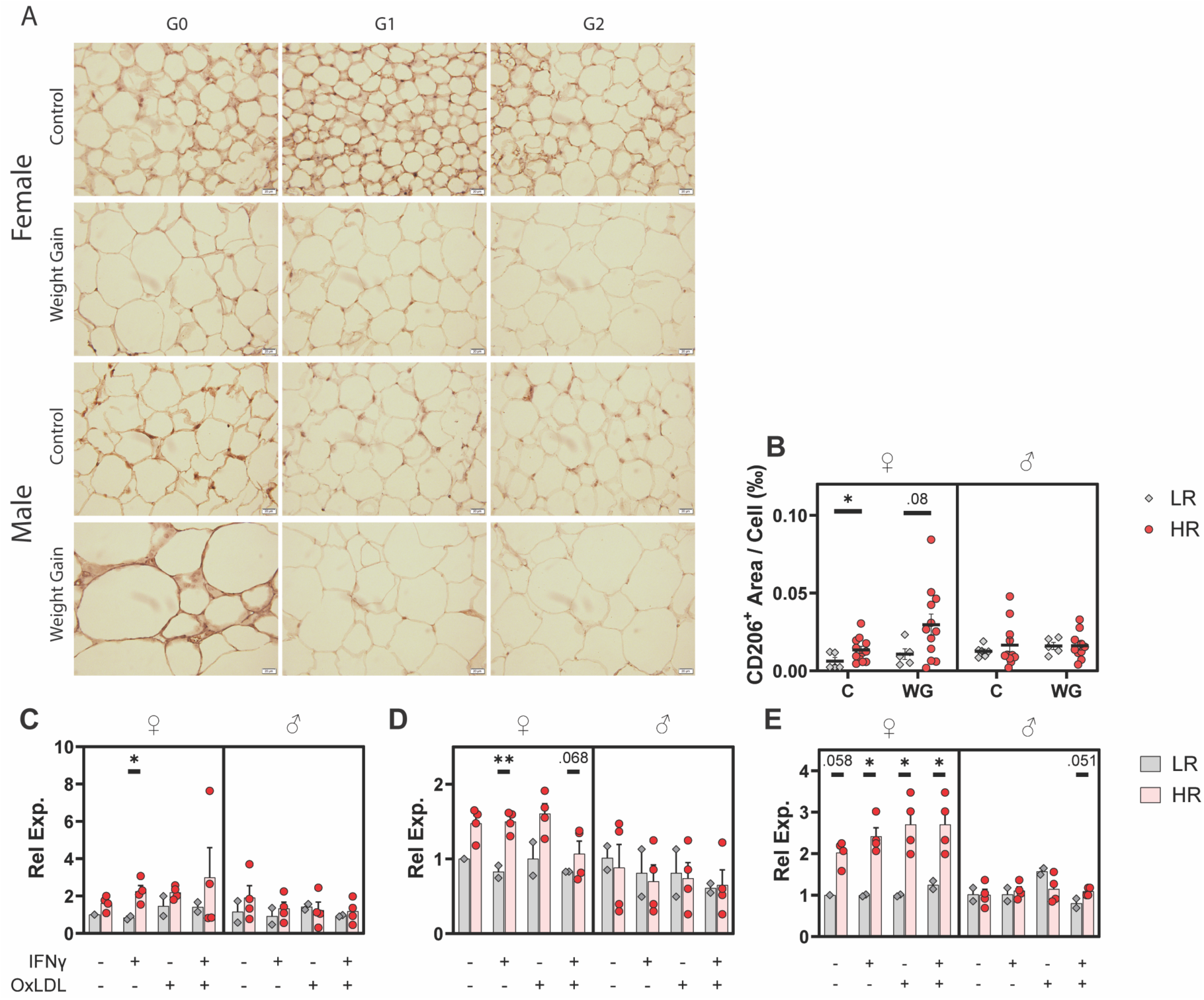
Immune Perturbations. **A)** Representative IHC images of iWAT stained for CD206. Scale bar, 20µM. **B)** Quantification of CD206-stained iWAT collected from short-term (16-week) experiment. CD206^+^ area was normalized to the number of cells in-frame. **C)** mRNA expression of *Abca1* in ApoL1+ BMDMs treated with OxLDL and/or IFNγ. Normalized to *Atpfa1*. **D)** mRNA expression of *Hmox1* in ApoL1+ BMDMs treated with OxLDL and/or IFN**γ**. **E)** mRNA expression of *Srebpf1* in ApoL1+ BMDMs treated with OxLDL and/or IFNγ. IHC, immunohistochemistry; iWAT, inguinal white adipose tissue; BMDM, bone marrow-derived macrophage; OxLDL, oxidized low-density lipoprotein. IFN**γ**, interferon-gamma

To further investigate the differences in macrophage phenotype based on *APOL1* genotype, we generated bone marrow derived macrophages (BMDMs) and exposed them to interferon gamma (IFNγ) and/or oxidized low-density lipoprotein (OxLDL), mimicking the hyperlipidemia often seen in metabolic syndrome, to induce inflammation. Parallel work we performed using this model system found increased inflammasome formation and decreased sensitivity to pro- resolving macrophages [47], prompting further gene-specific investigation. We found overexpression of the markers *Abca1*, *Hmox1*, and *Srebf1* in HR female macrophages, with differences in male macrophages being less pronounced to not present (Figures 4C-E). *Hmox1* expression in macrophages is important for the clearance of Hb-Hp and heme-Hx complexes, with failure to do so creating reactive oxygen species that are harmful to endothelial cells [48, 49]. *Abca1* expression moves cholesterol out of macrophages, which is important for preventing atherosclerotic foam cell formation [28]. *Srebf1* expression is mostly characterized in hepatocytes but has been shown to have some pro-inflammatory and some anti-inflammatory properties in macrophages [50, 51]. In aggregate, these results point towards a subtle sex-specific alteration in macrophage phenotype correlated with *APOL1* genotype. These results do not give reasoning for causality, however. Interestingly, Zhang et al. 2015 [52], from which the BMDM generation procedure used here is derived, observed their HR BMDMs to express lower levels of *Abca1* and have blunted cholesterol efflux. Their study looked exclusively at male macrophages, and as shown here they may be important sexual dimorphism to account for which may partially explain this disparity.

## DISCUSSION

In this study, we found that the deposition of fat mass in response to HFD differs between LR and HR *APOL1* mice in a sex-dependent manner. Specifically, female mice appear protected against DIO, but this protection is blunted by the presence of two HR *APOL1* alleles (Figure 1D, 1G). This observation phenocopies an association between weight and *APOL1* reported by Nadkarni et al. (2021) [36], with an additive increase in BMI of 0.36 kg/m^2/allele. When stratified by sex, they found a significant association between overweight risk, obesity risk, free fat mass index, and BMI in women, while only obesity risk proved significant in men. While this does not corroborate the inverse association we observed with our male mice, their finding provides some evidence that *APOL1* HR alleles may affect females more strongly than males. Together with our data, this may point towards a role in *APOL1* HR in increasing obesity odds in African American women [53].

There may be an evolutionary benefit provided by this elevated adiposity. Trypanosomiasis is known to alter the lipid profile of patients, with a marked increase in free triglycerides, cholesterol, and LDL, along with a drop in blood glucose [15, 54–57]. Trypanosomes are known to infiltrate adipose tissue, and the immune response to their presence is thought to be a key reason for the rapid fat loss patients experience [58]. Intriguingly, Machado et al. 2023 [58] found that lipolysis and the release of free fatty acids is toxic to adipocyte-resident trypanosomes, and blocking ATGL-lipolysis increased parasite load and decreased survival in mice. It is possible that the increased adiposity induced by *APOL1* HR genotype provides additional anti-trypanosome defense by promoting a larger store of fat that can be weaponized against parasites.

Additionally, an important distinction in assessing if adipose gain is potentially pathogenic is whether adipose tissue is growing by hypertrophy or hyperplasia. Hyperplasic growth occurs through the addition of new adipocytes without an increase in cell size. Hypertrophic growth occurs when adipocytes grow in size without increasing cell counts, and is associated with impaired angiogenesis, tissue hypoxia, insulin sensitivity, and low-grade inflammation [59, 60]. This low-grade inflammation is implicated in the development of cardiovascular comorbidities [45]. We identified hypertrophic growth in HR female mice after 16wk of HFD (Figure 2A-B), in accordance with their increased fat mass. This hypertrophic growth should drive higher levels of inflammation in the tissue. However, this was not readily apparent through transcriptional or histological analysis of adipose tissue, or markers of systemic inflammation analyzed in blood sera (data not presented). A possible explanation for this discrepancy is the crude nature of analyzing adipocyte size through histology compared to more robust methods [61]. Further characterization of adipocyte size in *APOL1* mice may prove fruitful, as definitive hypertrophy of adipocytes in *APOL1* HR mice may lend credence to the anti-trypanosome hypothesis detailed above.

Unfortunately, our other findings suggest that our model system could not appropriately induce a DIO phenotype in these mice. We did not see convincing evidence of hypertension (Figure 3A) in the absence of a high-salt diet, suggesting that HR APOL1 in endothelium is not sufficient for the hypertension phenotype. oGTTs confusingly showed that mice fed LFD had higher blood sugar levels, the opposite of what would be expected. Insulin tolerance testing data was not collected, although it likely would have demonstrated similar results. We also saw no difference in weight gain or fat mass for long-term WG mice (Figure 1G-H), despite both the short-term and long-term WG groups being on HFD for a total of 16 weeks prior to data collection. The two groups differ by the age at which HFD exposure began: 6 weeks postpartum for the short-term versus 16 weeks postpartum for the long-term. Other studies have looked at the effect of beginning HFD within different windows in mice. However, these studies tend to begin diet manipulation earlier than our study and find the critical window to be pre-or-peripubertal [62–64]. During this period, exposure to HFD appears to “prime” mice for health later in life. For instance, Cordoba-Chacon et al. (2012) [63] found that mice fed 16 weeks of HFD starting at 28 days postpartum had higher insulin sensitivity and higher lean mass than those started on HFD at 84 days postpartum. Our results here might point towards another critical window between 16-24 weeks postpartum in which APOL1-mediated DIO is most inducible by initiation of HFD, but this more likely reflects an inability of this system to replicate the results found.

We suspect the feeding behavior of the animals may have been influenced by their strain as well as their housing conditions, partially explaining these unexpected findings. FVB/NJ mice have historically been considered resistant to DIO, although Nascimento-Sales et al. (2017) [65] report that FVB/NJ mice are susceptible to DIO, and in fact become more insulin resistant and experience greater adipose inflammation than C57Bl/6J mice. These results were not replicated in our study. In order to circumvent these concerns and guarantee the development of obesity in a reliable and speedy fashion, we would recommend future researchers to backcross this *APOL1* expression system onto C57Bl/6J mice, speficially Lep^ob/ob^ or A^y^/a obesigenic systems [66].

Social dominance may have also affected individual eating habits. Social hierarchies in mice are known to influence eating behavior, with more dominant mice enjoying greater access to food [67]. Mice were housed three to a cage during these experiments, and the emergence of a dominant female was evident, particularly in WG cages (data not presented). However, it should be noted that single housing stresses animals through social isolation and an inability to huddle for warmth, factors which also alter food intake and weight gain. However, while expensive, single housing remains the standard for metabolic studies such as the one conducted here [67], and so our housing of multiple to a cage is a significant limitation.

In summary, this study provides limited evidence that *APOL1* high-risk genotypes may be involved in the pathogenesis of obesity in female but not male mice. This may provide important early insight into the disparity in obesity prevalence between female and male African Americans. However, the inability to produce a robust DIO-phenotype limits the applicability of these results. In particular, while we did see preliminary evidence of differences in macrophage and cardiac function, future models may benefit from inducing a stronger cardiovascular phenotype, including but not limited to hypertension and HFpEF [68]. However, we stress that a failure to distinguish sex would have missed the key finding reported here. This study most importantly highlights the importance of identifying sex as a relevant biological variable in the study of APOL1, both in future animal studies as well as human studies.

**Table 1.**
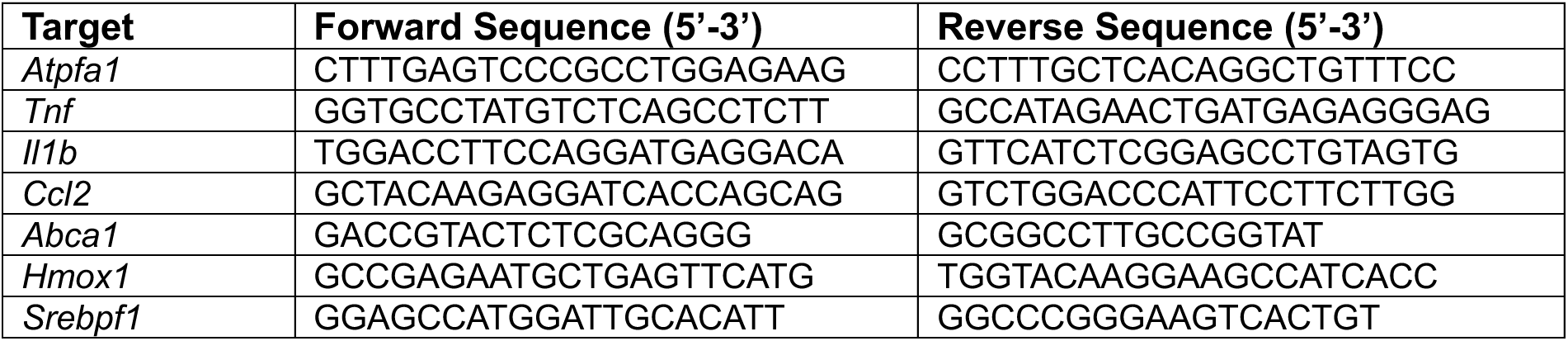
Primer sequences for RT-qPCR reactions.

## ACKNOWLEDGMENTS

EchoMRI and Luminex multiplexing services were provided by the Comprehensive Metabolic Core at Northwestern University.

Histology and immunohistochemistry services were provided by the Northwestern University Mouse Histology and Phenotyping Laboratory (MHPL) which is supported by the NCI P30-CA060553 grant awarded to the Robert H. Lurie Comprehensive Cancer Center.

A portion of the research reported in this publication was supported by the National Institute of Diabetes and Digestive and Kidney Diseases of the National Institutes of Health under Award Numbers U2CDK129917 and TL1DK132769. JL, MW, and AOK were supported by R01DK131521. The content is solely the responsibility of the authors and does not necessarily represent the official views of the National Institutes of Health.

## AUTHOR CONTRIBUTIONS

**Conceptualization:** JL

**Data Curation:** AK, JY, EL, MW

**Formal Analysis:** AK, MW, JY

**Investigation:** AK, JY, EL, MW, MC, JK

**Methodology:** JL, ET, GB

**Resources:** JL, ET, GB

**Supervision:** JL

**Visualization:** JL, AK

**Writing – original draft:** AK, MW

**Writing – reviewing & editing:** AK, JL, ET, GB, EL, MC, JK, JY, MW

## Supporting Information

**Supplemental Figure S1.**
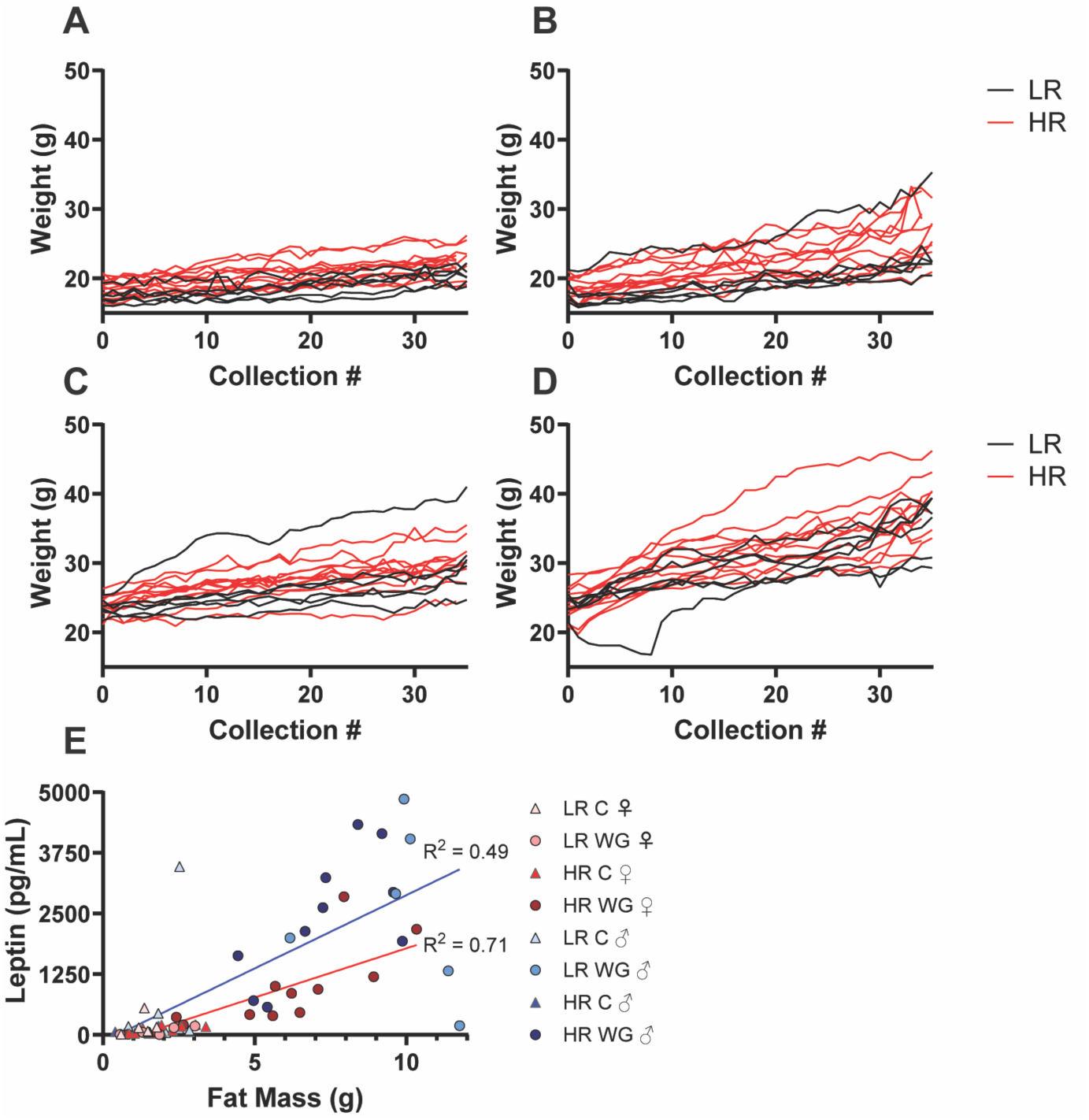
Individual gross weight-over-time of 16wk experiment. **A)** C Females **B)** WG Females **C)** C Males **D)** WG Males. Collection # increments per instance of being weighed. **E)** Correlation between fat mass and fasted serum leptin level. C, control; WG, weight gain.

**Supplemental Figure S2.**
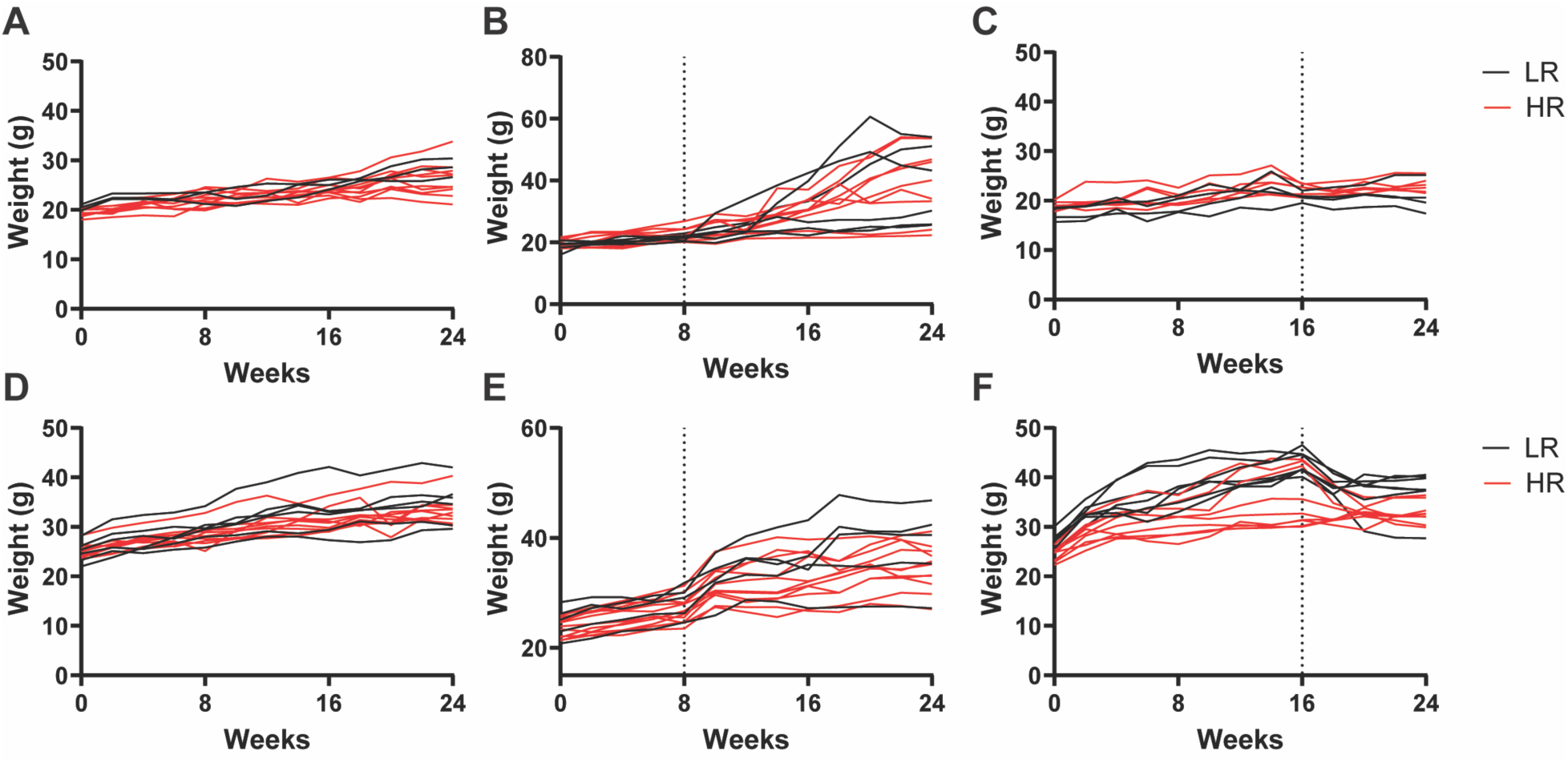
Individual gross weight-over-time plots of 24wk experiment. **A)** C Females **B)** WG Females **C)** WL Females **D)** C Males **E)** WG males **F)** WL males. C, control; WG, weight gain; WL, weight loss.

**Supplemental Figure S3.**
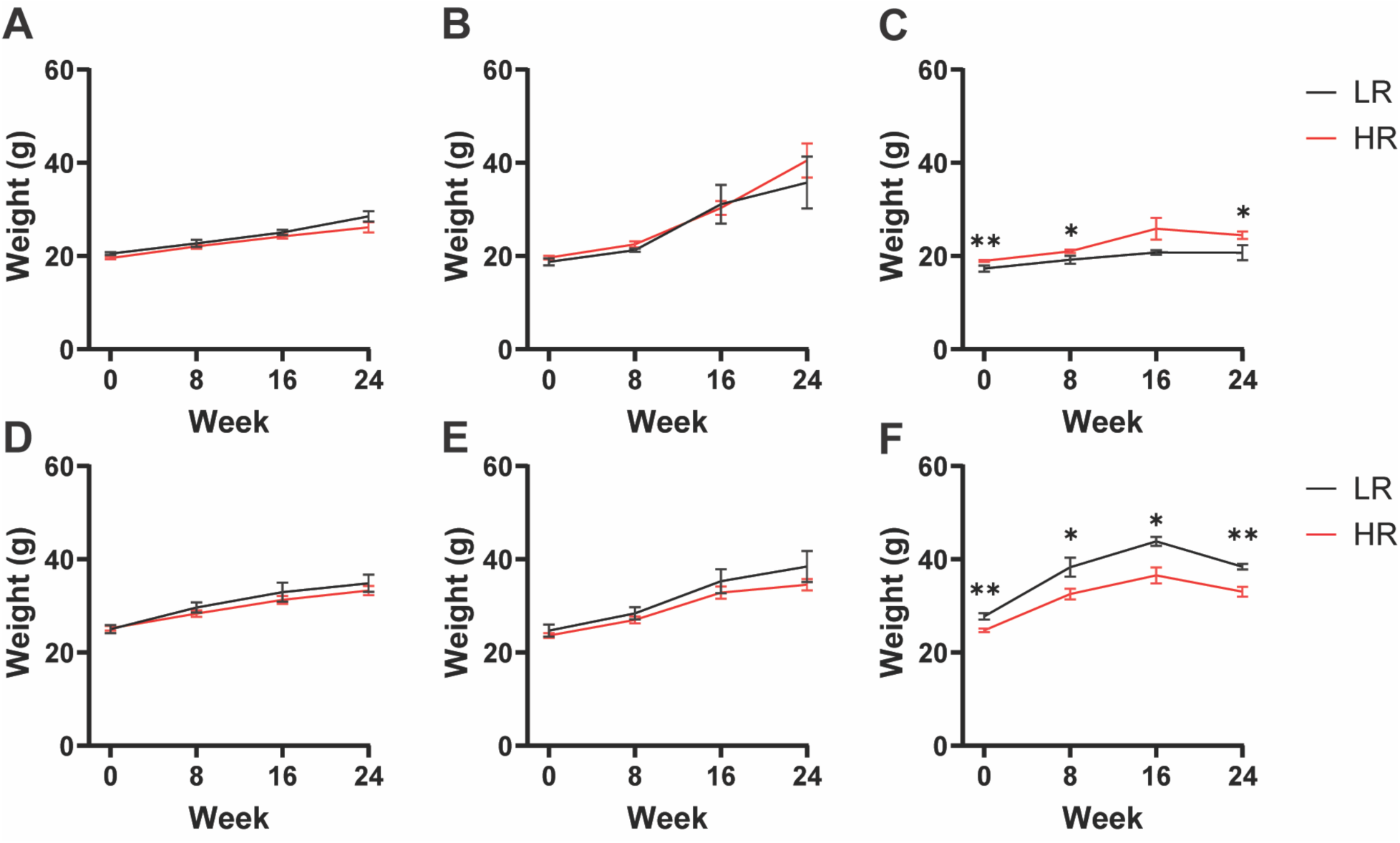
Gross weight at 8, 16, and 24 weeks of 24wk experiment. **A)** C Females **B)** WG Females **C)** WL Females **D)** C Males **E)** WG males **F)** WL males. C, control; WG, weight gain; WL, weight loss.

**Supplemental Figure S4.**
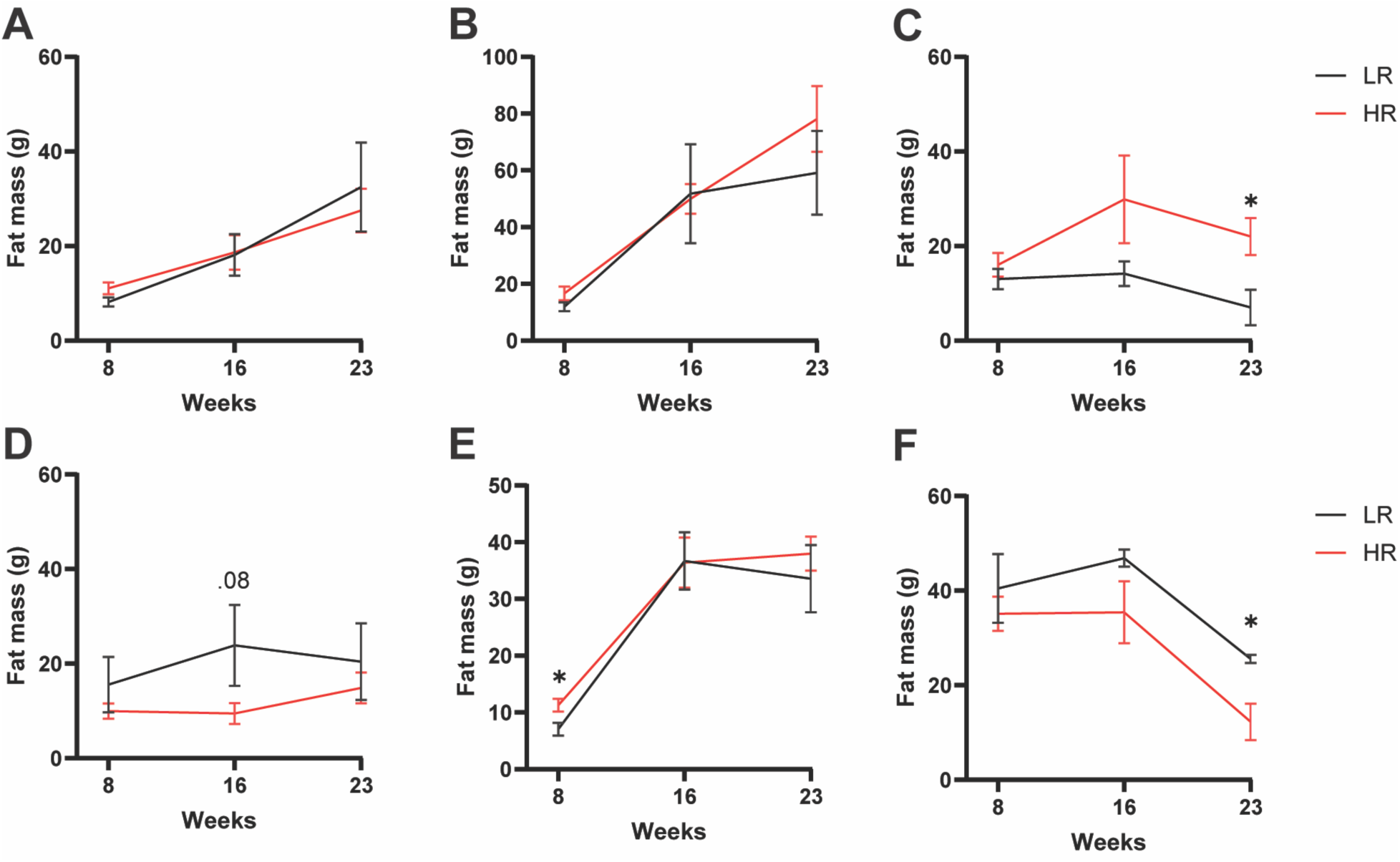
Fat mass at 8, 16, and 24 weeks of 24wk experiment. **A)** C Females **B)** WG Females **C)** WL Females **D)** C Males **E)** WG males **F)** WL males. C, control; WG, weight gain; WL, weight loss.

**Supplemental Figure S5.**
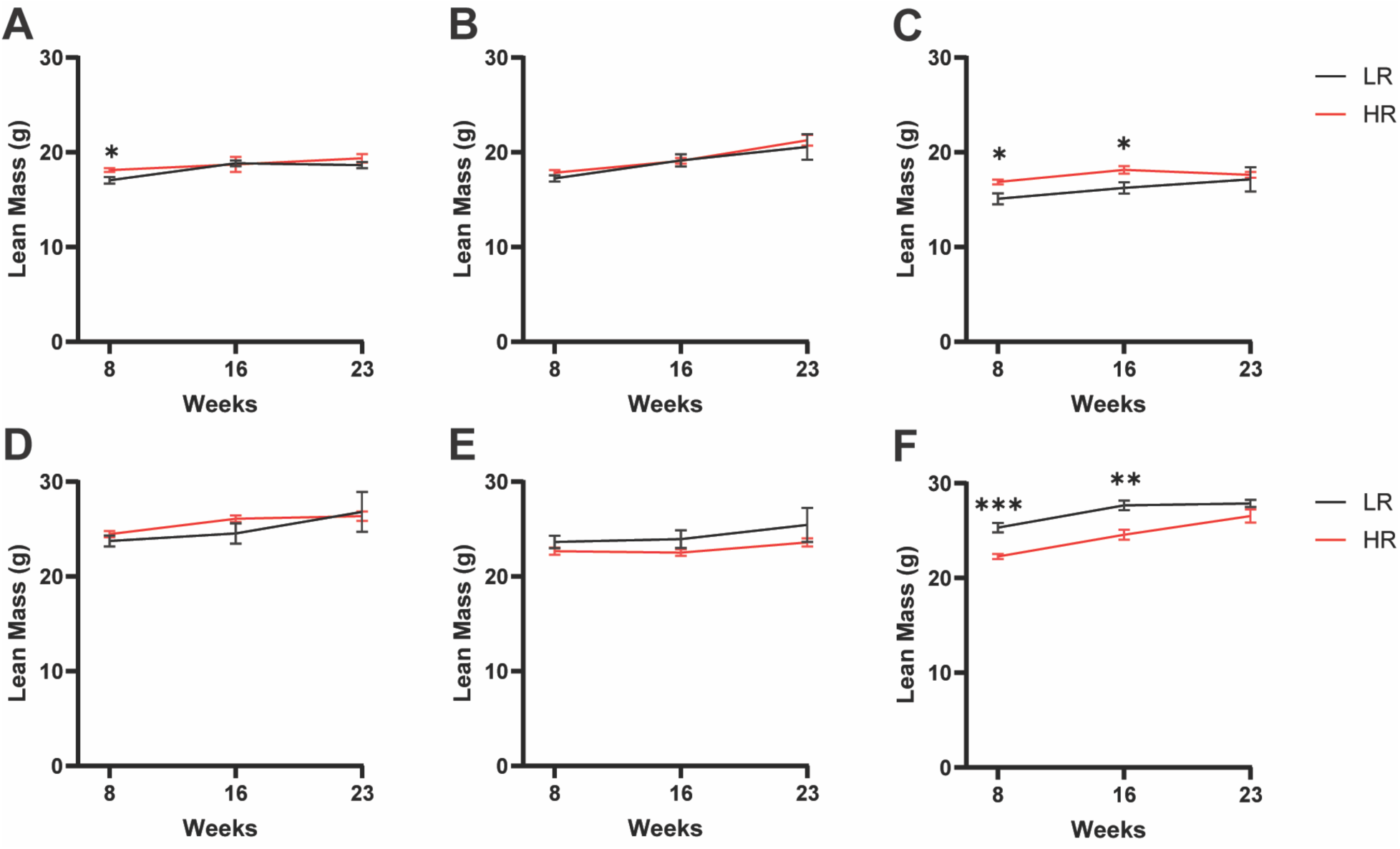
Lean mass at 8, 16, and 24 weeks of 24wk experiment. **A)** C Females **B)** WG Females **C)** WL Females **D)** C Males **E)** WG males **F)** WL males. C, control; WG, weight gain; WL, weight loss.

**Supplemental Figure S6.**
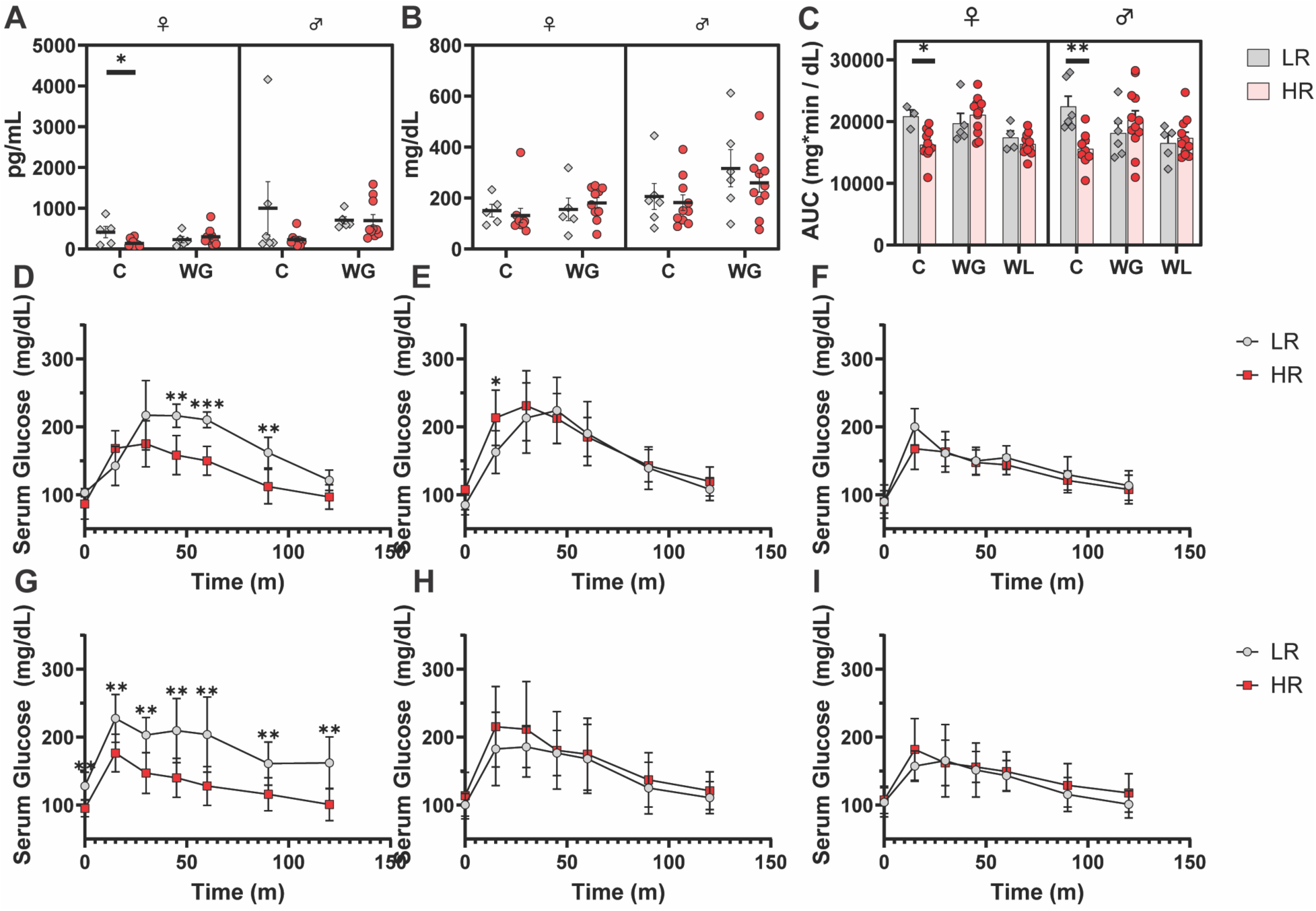
oGTT plots. **A)** Fasted serum insulin concentration of 16wk experiment. **B)** Fasted serum glucose concentration of 16wk experiment. **C)** Area-under-the-curve of 24wk experiment oGTTs. oGTT curves of 24wk **D)** C Females **E)** WG Females **F)** WL Females **G)** C Males **H)** WG males **I)** WL males. oGTT, oral glucose tolerance test; C, control; WG, weight gain; WL, weight loss.

**Supplemental Figure S7.**
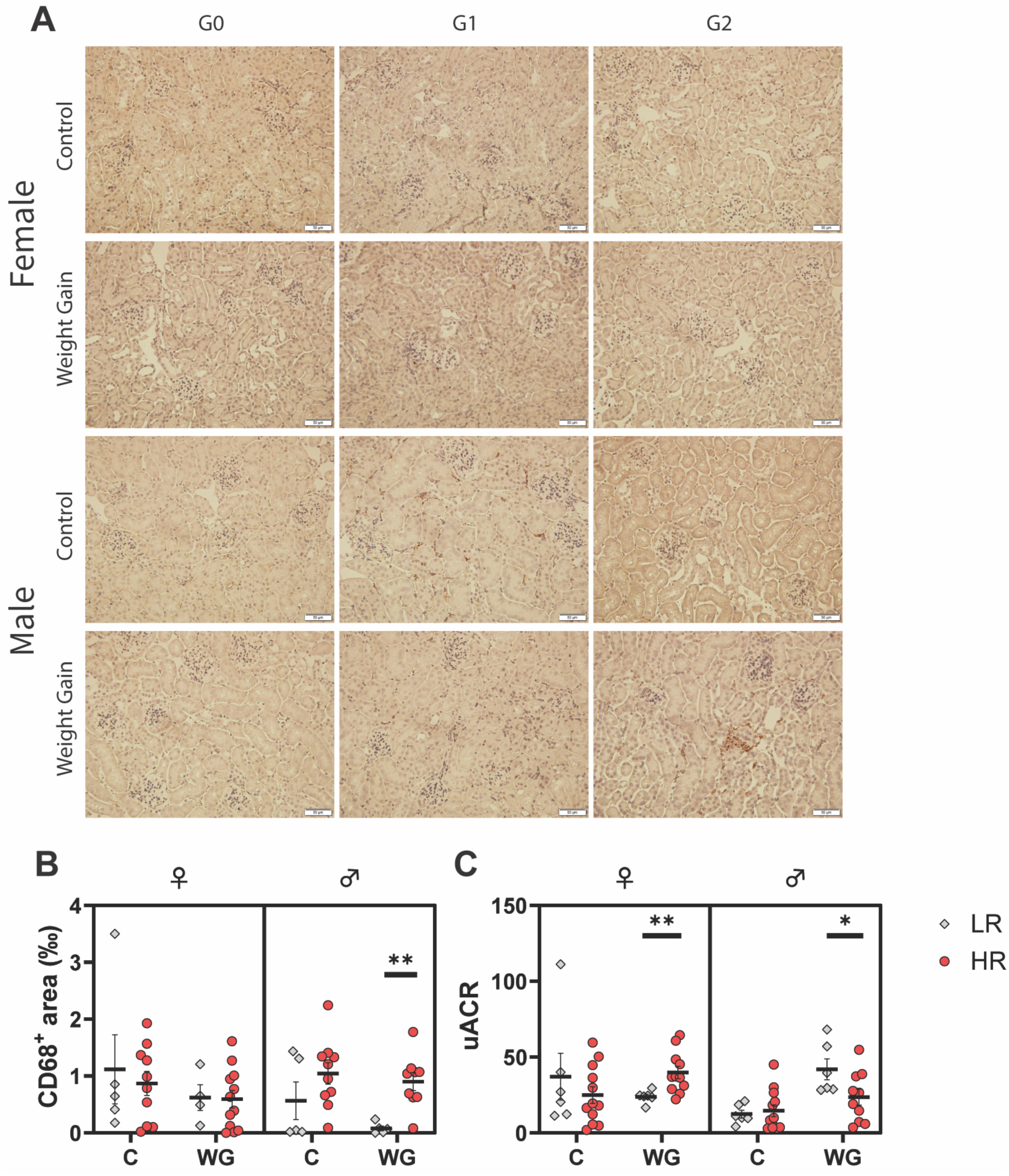
Kidney Injury. **A)** Representative IHC images of kidney bisections stained for CD68. Scale bar, 20µM. **B)** Quantification of CD68-stained kidney bisections collected from short-term (16-week) experiment. **C)** uACR of 16wk experiment. IHC, immunohistochemistry; uACR, urinary albumin:creatinine ratio.

